# Interoception vs. Exteroception: Cardiac interoception competes with tactile perception, yet also facilitates self-relevance encoding

**DOI:** 10.1101/2025.06.25.660685

**Authors:** Marie Loescher, Patrick Haggard, Catherine Tallon-Baudry

## Abstract

Internal bodily signals, notably the heartbeat, influence our perception of the external world – but the nature of this influence remains unclear. Different frameworks, originating in opposing views of the function of interoception, have developed largely in parallel. One line of evidence (Internal/External Competition) indicates that interoceptive and exteroceptive inputs compete for neural resources. Another line (Self-related Facilitation) shows a link between interoceptive and self-related processing, which might include computing the self-relevance of exteroceptive inputs. We contrasted these accounts within a single experimental task for which they yielded distinct predictions. We measured heartbeat-evoked potentials (HEPs, a measure of cardiac interoception) with EEG, and manipulated the self-relevance of an audio-tactile stimulus by placing the audio source either inside or outside the peripersonal space immediately around the body. On one hand, pre-stimulus HEP amplitudes over somatosensory cortex were linked to slower reaction times, and affected audio-tactile stimulus-evoked responses in the same area, indicating competition for shared neural resources. On the other hand, pre-stimulus HEPs over integrative sensorimotor and default-mode network regions facilitated stimulus self-relevance encoding, both in reaction times and audio-tactile evoked responses. Importantly, Competition and Facilitation effects were spatially and statistically independent from each other. We therefore reconcile the two views by showing the co-existence of two independent mechanisms: one that allocates neural resources to either internal bodily signals or the external world, and another by which interoception and exteroception are combined to determine the self-relevance of external signals. Our results highlight the multi-dimensionality of HEPs, and of internal states more generally.

**Significance Statement:** Do internal bodily signals distract us from the external world (Internal/External Competition account)? Or do internal signals contribute to conscious perception, by situating the perceived external world relative to the organism (Self-related Facilitation account)? So far, both accounts – reflecting fundamentally different views of brain-body interactions – received experimental support, but have never been compared directly. We measured neural responses to heartbeats, and tested how they influenced perception in an audio-tactile reaction time task where self-relevance was manipulated. We found evidence for both accounts, reflecting independent mechanisms in distinct brain regions. Our results reconcile two hitherto independent and seemingly contradictory research programmes on the relationship between interoception and exteroception. They further highlight the multi-dimensionality of cardiac-brain interactions, and hence of internal state.

## Introduction

Although we often think of the brain as primarily responding to environmental (*exteroceptive*) inputs, a large part of its activity is also dedicated to processing *interoceptive* information from internal organs (1). For example, each heartbeat elicits a wide-spread neural response, heartbeat-evoked potentials (HEPs), even in areas traditionally associated with exteroception such as somatosensory cortex (for review, see 2). How, then, does such – largely unconscious – interoceptive processing interact with exteroceptive perception? The burgeoning literature on this question appears divided between a “Internal/External Competition” account, according to which interoceptive and exteroceptive inputs compete for neural or attentional resources, and a “Self-related Facilitation” account, where interoceptive signals support subjective aspects of exteroception. These frameworks reflect a deeper divide on the function of brain-body interactions: while for one side, the processing of interoceptive signals primarily serves to regulate the body’s physiological state, the other side assigns to bodily signals functions far beyond autonomic regulation, contributing even to the generation of a self. Competition and Self-related Facilitation have so far developed in parallel, each guiding largely independent lines of research. In the present work, we experimentally test these two opposing accounts.

According to the Internal/External Competition view, interoceptive inputs inform the brain about bodily states much like exteroception informs about the world. Given finite resources, the brain would prioritise the processing of either interoceptive or exteroceptive information, in the same way as multiple external inputs can compete for attention. Top-down attention to heartbeats is known to increase HEP amplitude (3–5). By extension, spontaneous fluctuations in HEPs have sometimes been interpreted as reflecting spontaneous attentional shifts between interoception and exteroception (e.g., 5–7). In particular, this interpretation was put forward to explain why large pre-stimulus HEP amplitudes are associated with reduced detection of near-threshold tactile stimuli (6, 8). Competition for shared resources is especially plausible between cardiac *vs*. tactile processing: the primary somatosensory cortex is a known neural source of HEPs and therefore a possible site of competition (8–10), and somatosensory pathways also convey cardiac afferences (11–13).

The Self-related Facilitation account, by contrast, holds that bodily signals may tend to enhance exteroception, in particular its subjective aspects (e.g., 14), rather than compete with it. On this view, interoceptive afferents not only inform about the state of an organ, but also contribute to a neural representation of the organism as a unified entity, i.e., to an embodied “self” that can feel (15–18) – a simple form of self that does not require an explicit, metacognitive self-representation. Experimentally, HEPs have been linked to such simple forms of self across cognitive domains, such as in emotion (19) imagination (20), body representation (21), and spontaneous thought (22). Consequently, cardiac interoception may also facilitate self-related aspects of exteroception, such as appraising the self-relevance of an external stimulus. Such an interpretation was notably proposed by Park and colleagues (14), who found that higher HEP amplitudes predicted better detection of a near-threshold visual stimulus and indexed a self-related process, since perception at threshold is intrinsically subjective (i.e., self-related). However, the experiment did not directly manipulate a self-related dimension of exteroception.

The Internal/External Competition and Self-related Facilitation accounts thus make opposing predictions, respectively suggesting a positive or negative relation between HEP amplitude and exteroception. We aimed to test these predictions within the same task, by including elements of both self-related processing and environmental perception. Specifically, we manipulated the self-relevance of a tactile stimulus using an auditory modulation of peripersonal space (PPS), the area of external space immediately surrounding the body. While initially defined by distance alone (23), PPS is now understood as a space of enhanced behavioural relevance, tightly linked to a bodily sense of self (24–26), as it also depends on factors like stimulus valence or the subjects’ own body size and motion (27–29). This conceptualisation of PPS emphasizes that self-relevance does not always have to involve explicit self-reference, such as a learned association to oneself (e.g. in 30). Rather, self-relevance here simply involves the encoding of stimuli *relative to* ourselves: to our location in space, but also our current physiological needs and so forth.

We manipulated the self-relevance of a tactile stimulus by presenting simultaneous tones either within or beyond PPS (31), while recording both electroencephalogram (EEG) and electrocardiogram (ECG) (fig.1a-c). Tactile stimuli accompanied by near sounds are perceived as more relevant, resulting in faster tactile reaction times than those accompanied by distant sounds – we term this difference the *self-relevance effect*. Internal/External Competition and Self-related Facilitation produce distinct predictions for this task: If interoception *competes* with exteroception, stronger HEPs in the time-window preceding stimulation should show a general association with longer subsequent reaction times (fig. 1d). If interoception *facilitates* specifically self-related aspects of exteroception, stronger pre-stimulus HEPs should amplify the self-relevance effect (yielding faster tactile responses with sounds in near space, relative to far space) – in other words, HEPs should specifically interact with peripersonal space (fig. 1e).

**Figure 1.**
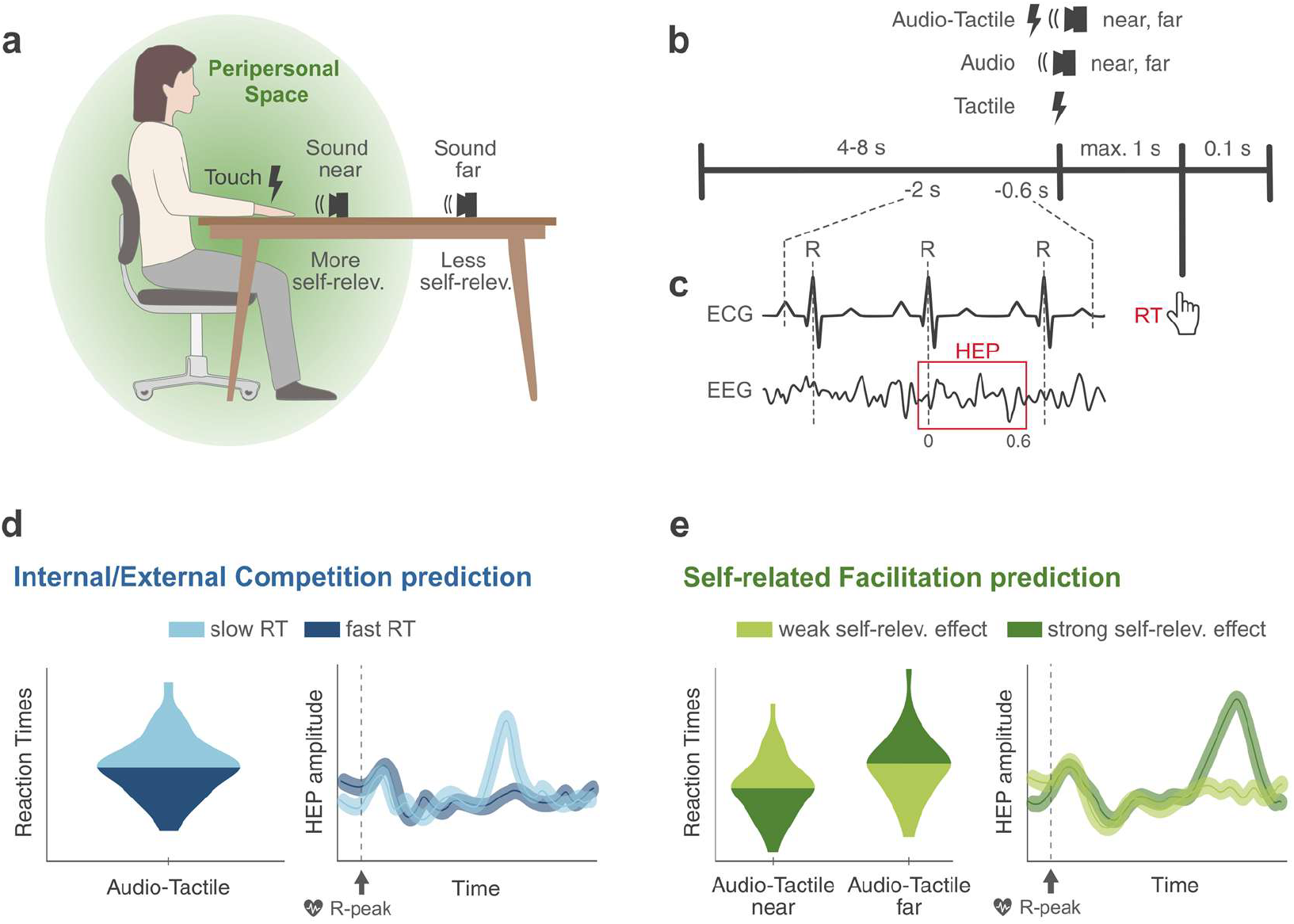
Experimental paradigm. **a)** Experimental setup. 37 participants responded to non-painful electrical stimuli on their hand. The self-relevance of the tactile stimulus was manipulated using the subject’s peripersonal space, within which stimuli are perceived as more relevant: tactile stimuli could be accompanied by a short sound played either from a speaker close to the subject’s hand, or from a speaker placed at 130cm distance. **b)** Trial structure. After a jittered delay, one of five different stimulus conditions could be presented (Audio-Tactile near, Audio-Tactile far, catch conditions Audio near and Audio far, and control condition Tactile). Participants responded as fast as possible to tactile stimuli (i.e., AT or T trials), while ignoring the sounds. **c)** Measures. Besides RT, we also recorded the ECG and EEG. We measured the HEP to the last heartbeat before stimulus onset. **d)** Predictions of the Internal/External Competition account. Slower RT (lighter blue) should be preceded by higher pre-stimulus HEP amplitudes. **e)** Predictions of the Self-related Facilitation account. Higher HEP amplitudes should correlate with enhanced self-relevance processing (darker green), i.e., faster RT for more self-relevant stimuli (in ATnear), and slower RT for less self-relevant stimuli (in ATfar). Abbreviations: ECG: Electrocardiogram, EEG: Electroencephalogram, RT: Reaction Times, HEP: Heartbeat-Evoked Potential, ATnear: Audio-Tactile near, ATfar: Audio-Tactile far.

## Results

### 1. Behavioural results

37 participants detected non-painful electrical stimuli on their hand, while ignoring concurrent sounds when they were played. On average, participants detected 96.53 ± 4.04 % (mean ± SD; range: 85-100 %) of all tactile stimuli with a false alarm rate of 1.98 ± 1.78 % (range: 0-7.50 %) and a grand average reaction time of 440 ± 72 ms (range of means: 281-653 ms).

#### Faster reaction times for more self-relevant tactile stimuli

We first confirmed the presence of a self-relevance effect, i.e., an effect of sound distance on tactile reaction times (31), using a 2×2 repeated-measures ANOVA on the log-transformed reaction times of audio-tactile (AT) trials with both sound distance condition (ATnear: touch-concurrent sound was played close to the stimulated hand, or ATfar: sound was played far from the hand) and cardiac phase at the time of stimulation (systole, diastole) as within-subject factors. Sound distance significantly affected tactile reaction times (F(1,36) = 9.49, p = 0.004; untransformed mean ± SD reaction times of ATnear = 425 ± 74 ms; of ATfar = 431 ± 73 ms). Whether the stimulus occurred during cardiac systole or diastole did not have any significant effect on reaction times (F(1,36) = 2.54, p = 0.120). The interaction of cardiac phase and sound distance did not reach significance (F(1,36) = 3.30, p = 0.077). Overall, we thus successfully replicated the paradigm previously validated by Ronga et al. (31): By playing a concurrent sound either within or outside of participant’s peri-personal space, we manipulated the relevance of the tactile stimulus for the participant, reflected in shorter reaction times for ATnear compared to ATfar trials.

#### No pre-stimulus difference in arousal

We verified that the self-relevance effect could not be explained by a difference between ATnear and ATfar trials in pre-stimulus arousal state, measured with cardiac inter-beat intervals and pupil diameter. We found no significant difference and moderate evidence for an absence of difference in arousal between ATnear and and ATfar trials, both in the average inter-beat intervals of the three heartbeats preceding stimulation onset (two-tailed paired t-test, t(32) = 0.21, p = 0.835, Bayes Factor in favour of correlation (BF10) = 0.19), or in average pupil diameter across the second preceding stimulation onset (t(32) = −0.58, p = 0.567, BF10 = 0.22).

### 2. Pre-stimulus heartbeat-evoked potentials predict both tactile and self-relevance processing

We first removed the component of reaction times related to experimental block order and cardiac parameters (inter-beat interval and cardiac phase at the time of stimulation) using a first general linear model (GLM) (Eq.1). We used the residual Reaction Times (RT_res_) for all subsequent analyses. We then tested the Competition and the Self-related Facilitation accounts’ predictions by statistically modelling pre-stimulus HEPs with both RT_res_ and the interaction between RT_res_ and near vs. far sound distance condition (GLM in Eq.2). In other words, we tested where and when the pre-stimulus HEP predicted subsequent reaction times and/or the effect of sound distance on reaction times (i.e., the self-relevance effect). Note that sound distance is not included as a main regressor because pre-stimulus HEPs could not predict future stimulus condition on its own. For each trial, the HEP was defined as the last 600 ms epoch following a cardiac R-peak and ending before stimulation onset, such that the entire HEP epoch always preceded the stimulus. According to the Internal/External Competition account, the neural processing of heartbeats should compete with the processing of exteroceptive stimuli. Consequently, larger HEP amplitudes should be associated with slower reaction times (fig. 1d). The Self-related Facilitation account proposes that the neural processing of heartbeats supports self-related aspects of exteroceptive perception. Accordingly, trials with large pre-stimulus HEP amplitudes should also be trials with a strong self-relevance effect (fig. 1e), corresponding to an interaction effect between reaction times and sound distance in the GLM.

#### Competition and facilitation co-exist: HEP in posterior somatosensory cortices compete with tactile inputs, HEPs in prefrontal, temporal and premotor cortices facilitate self-relevance processing

Main results of the HEP analysis are summarised in figure 2. We found that higher pre-stimulus HEP amplitude between 454 and 494 ms after the R-peak (cluster sum(t) = 850.29, Monte-Carlo *p* = 0.026) in a right central cluster (11 electrodes: Pz, TP8, CP6, CP4, CP2, P2, P4, P6, P8, PO8 and PO4) were associated with slower reaction times, i.e., the signature of a competition between tactile processing and cardiac processing (fig. 2a-c). Source localisation analysis revealed that the anatomical region with the largest contribution to this Competition effect (fig. 2d) was in the right postcentral sulcus, in posterior somatosensory cortex (MNI(x,y,z) = 50 −22 40).

**Figure 2.**
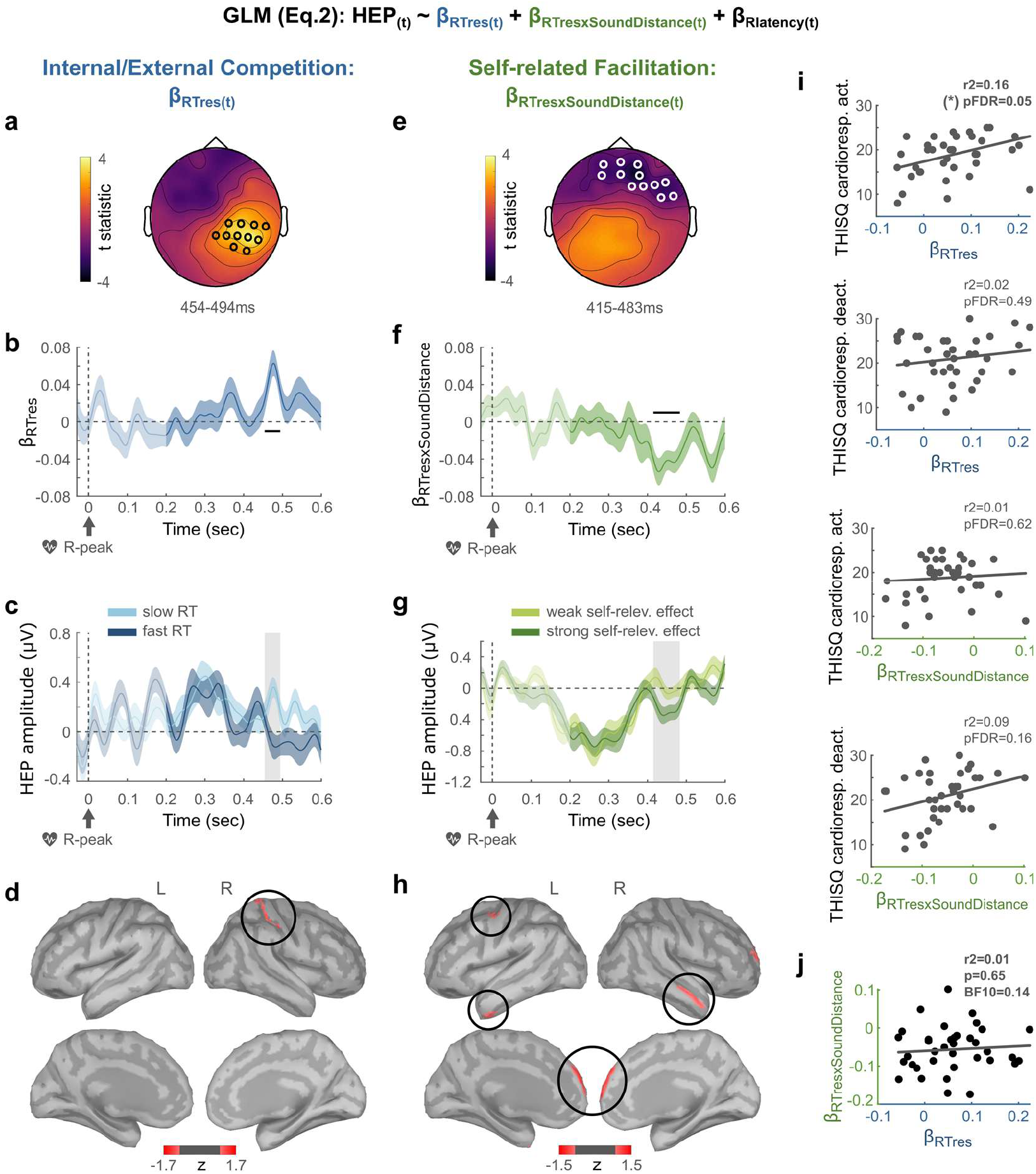
Pre-stimulus HEPs compete with tactile processing (a-d), but also facilitate self-relevance encoding (e-h). **a)** Topography of the cluster during which there was an Internal/External Competition effect, i.e., a significant link between pre-stimulus HEPs and tactile reaction times (RT_res_), averaged over the significant time-window. **b)** Timecourse of the estimate β_RTres_ averaged over the significant channels highlighted in (a). Shaded area depicts standard error. Lighter colour indicates un-tested time-window. The black bar indicates the significant time-window. **c)** For visualisation purposes only, HEPs in significant channels in (a) were averaged across the 50 % of trials with slower RT (light blue) vs. the 50 % of trials with faster RT (dark blue). During the time-window of the significant cluster for β_RTres_ (grey area), HEP amplitudes were higher for slow RT trials. **d)** The brain region mostly contributing to the Competition effect was localized in the right posterior somatosensory cortex (threshold at 54 % of max. z-value in a minimum of 20 adjacent vertices). **e)** Topography and **f)** time-course of the estimate β_RTresxSoundDistance_ in the cluster during which there was a significant Self-related Facilitation effect, i.e., a link between pre-stimulus HEPs and the self-relevance effect on RT. **g)** For visualisation purposes only, HEPs in significant channels in (e) were averaged across trials with a strong self-relevance effect (i.e., the fastest 50 % of ATnear trials, and the slowest 50 % of ATfar trials; dark green) versus trials with a weaker self-relevance effect (i.e., remaining trials; light green). During the time-window of the significant cluster for β_RTresxSoundDistance_ (grey area), absolute HEP amplitudes were higher for strong self-relevance effect trials. **h)** Brain regions mostly contributing to the Facilitation effect were localized in bilateral anterior temporal regions, left dorsal premotor cortex, and bilateral rostral prefrontal cortices. **i)** Correlations between HEP results and scores of self-reported interoceptive sensations with THISQ cardio-respiratory activation and deactivation subscales (35). The only link surviving multiple comparison correction is between the Competition effect and cardio-respiratory activation. **j)** Moderate Bayesian evidence for an absence of correlation between Competition and Facilitation HEP effects, across participants. Results for the control regressor β_Rlatency_ of the GLM (Eq.2), capturing HEP variance due to the timing of the R-peak relative to stimulus onset, are depicted in the supplementary fig. 1. **Abbreviations:** HEP: Heartbeat-Evoked Potential, RT: Reaction Time, BF10: Bayes Factor in favour of a correlation, pFDR: False-Discovery Rate corrected p-value.

We also found significant evidence for a Self-related Facilitation effect, beginning at slightly earlier latencies and at markedly distinct locations. Facilitation appears as a significant interaction effect of RT_res_xSoundDistance between 415 and 483 ms after the R-peak (cluster sum(t) = −1075.21, Monte-Carlo *p* = 0.014) in a right frontal cluster (12 electrodes: Fp1, AF3, Fpz, Fp2, AF4, AFz, F2, F4, F6, F8, FT8 and FC6) (fig. 2e-f). In particular, higher pre-stimulus HEP amplitudes over this cluster were associated with a stronger self-relevance effect (fig. 2g). The anatomical regions mostly contributing to this Facilitation effect were located in the bilateral rostral prefrontal cortices (MNI(x,y,z) = 9, 70, 13 and −6, 67, 21, respectively) and anterior temporal regions (MNI(x,y,z) = −46 −4 −34 and 59 7 −20, respectively), as well as the left dorsal premotor cortex (precentral sulcus, MNI(x,y,z) = −37, −5, 57) (fig. 2h).

Our results thus provide evidence relevant to both the Competition and the Self-related Facilitation accounts, but potentially linked to distinct aspects of cardiac interoceptive processing. We find that neural responses to heartbeats influence behaviour in two spatially (and, to some extent, temporally) distinct clusters. In what we will call the “HEP-Competition” cluster, larger responses predict slower reaction times, and in the “HEP-Facilitation” cluster, larger responses enhance the behavioural expression of stimulus self-relevance.

To verify that the observed Competition effect concerned tactile processing generally, and not only bimodal audio-tactile trials, we furthermore tested for the presence of a Competition effect in unimodal tactile (T) trials, using a separate GLM (Eq.3) on the times and electrodes of the HEP-Competition cluster. In T trials as well, higher HEP amplitudes were associated with slower reaction times (from 467 to 484 ms after the R-peak, over 9 electrodes: Pz, CP6, CP4, P2, P4, P6, P8, PO8 and PO4; cluster sum(t) = 307.68, Monte-Carlo *p* = 0.005).

#### Control analyses: No effects in ECG nor in permuted HEP data

To verify that the observed effects were due to neural HEPs, and not to a difference in the electrocardiogram itself, we applied the same GLM (Eq.2) to the ECG segments corresponding to the heartbeats used to compute HEPs. None of the three ECG derivations showed an effect of either RT_res_ or RT_res_xSoundDistance (no candidate clusters for RT_res_; 3 candidate clusters for RT_res_xSoundDistance, all with Monte-Carlo p ≥ 0.2). We can thus conclude that the effects are truly of neural origin rather than reflecting a difference in cardiac input (32).

Furthermore, pre-stimulus EEG often contains expectancy-related slow drifts ramping up to each stimulus (1, 33) which could affect mean HEP amplitudes depending on their timing relative to stimulation onset and lead to an effect measurable in, but not specific to, HEPs. To control for this confound, our GLM (Eq.2) also contained the timing of the HEP relative to stimulation onset as a third regressor. We report results pertaining to β_Rlatency(t)_ in the supplementary materials, showing that this regressor successfully captures variance in HEPs related to heart-unspecific slow drifts in the EEG (supplementary fig. 1). Second, we also verified that both Competition and Facilitation effects were time-locked to heartbeats using a permuted heartbeats analysis (RT_res_ Monte-Carlo p = 0.050; RT_res_xSoundDistance Monte-Carlo p = 0.014), effectively ruling out the possibility that our results could be entirely due to an underlying slow wave (1, 33).

#### Competition and Self-related Facilitation effects are independent from each other

Do the Competition and Facilitation effects reflect distinct underlying mechanisms, or is there a single underlying dimension such that strong Competition effects correlate with weak Facilitation effects and vice-versa? By design, the GLM terms on which the two effects are based are different: relation with RT for Competition, but interaction between sound distance and RT for Self-related Facilitation. The Variance Inflation Factors (VIFs) between regressors were small (all individual VIFs between any regressors < 1.34). We furthermore tested across participants for correlation between the two effects, and found moderate evidence for the absence of correlation (fig. 2j; robust correlation, r^2^<0.01, *p* = 0.65, BF10 = 0.14). We conclude that the effects are independent from each other.

#### The Competition effect, but not the Facilitation effect, is marginally linked to cardiac interoception scores

Lastly, we explored whether the HEP effects on behaviour were modulated by participants’ trait anxiety, quantified by STAI questionnaire scores (34), or by their sensitivity to cardiac sensations in everyday life, quantified by the cardiorespiratory activation and deactivation subscale scores of the THISQ questionnaire (35) (fig. 2i). We found moderate evidence for the absence of correlations between either of the two HEP effects and trait anxiety (Competition: r^2^ = 0.003, uncorrected p = 0.73, BF10 = 0.14; Facilitation: r^2^ = 0.016, p = 0.46, BF10 = 0.17), and moderate and weak evidence, respectively, for the absence of correlation with the cardiorespiratory deactivation subscale (Competition: r^2^ = 0.023, uncorrected p = 0.37, BF10 = 0.19; Facilitation: r^2^ = 0.085, p = 0.079, pFDR = 0.159, BF10 = 0.59, False-Discovery Rate (FDR) correction across the two beta scores and the two subscales). Interestingly, there was positive correlation between the Competition effect and the cardiorespiratory activation subscale that appeared as a trend after correction for multiple comparisons (r^2^ = 0.162, p = 0.013, pFDR = 0.054, BF10 = 2.66), but not for the Facilitation effect (r^2^ = 0.007, p = 0.624, BF10 = 0.14). The cardiorespiratory subscales contain separate questions pertaining to cardiac or to respiratory sensations. To explore whether the observed correlation could be attributed more to one or the other type of questions, we recomputed correlations between the Competition effect and the two question types separately. We observed a correlation between Competition and the cardiac activation questions (r^2^ = 0.203, uncorrected p = 0.005), but not the respiratory activation questions (r^2^ = 0.044, uncorrected p = 0.212). The link between the Competition effect and the cardiorespiratory activation scale therefore seems to be specifically related to cardiac interoceptive sensitivity.

### 3. Competing and Facilitatory HEP components differentially impact stimulus-evoked responses

We show that pre-stimulus HEP amplitudes over two distinct latencies and electrode sites (HEP-Competition and HEP-Facilitation) differentially affect behavioural responses. It follows that there should be two corresponding modulations of the neural stimulus-evoked response (SEP) to audio-tactile stimuli. How do exteroceptive and interoceptive responses interact? More precisely, when and where is the SEP affected by competition with cardiac processing, and when and where do we observe in SEP a self-related facilitation, i.e., an effect of the interaction between HEPs and self-relevance processing? We addressed these questions in a follow-up exploratory analysis. For each trial, we computed average pre-stimulus HEP amplitudes separately over the timepoints and electrodes of the Competition cluster (yielding HEP-Competition) and of the Facilitation cluster (yielding HEP-Facilitation). Competition and Self-related Facilitation accounts again produce distinct predictions: According to Competition, pre-stimulus HEP amplitude should negatively affect SEP amplitude. According to Facilitation, HEPs should modulate the effect of stimulus self-relevance on the SEP, corresponding to a HEP-FacilitationxSoundDistance interaction effect. To test these predictions, we modelled trial-by-trial SEPs in a GLM (Eq.4) using sound distance condition, HEP-Competition, HEP-Facilitation, and the interaction between HEP-Facilitation and sound distance as regressors.

Firstly, we verified that sound distance modulated the SEP of audio-tactile trials as expected from the literature (31), which we report in the supplementary materials (supplementary fig. 2). Then, confirming the Competition account’s prediction, we found a main effect of HEP-Competition on the SEP in two clusters: a positive posterior right cluster between 177 and 245 ms after stimulus onset (cluster sum(t) = 1300.72, Monte-Carlo *p* = 0.004, 14 electrodes: PO7, PO3, Iz, Oz, POz, C6, TP8, CP6, P4, P6, P8, P10, PO8, O2), and a negative frontal left cluster between 209 and 250 ms after stimulus onset (cluster sum(t) = −959.09, Monte-Carlo *p* = 0.015, 11 electrodes: F1, F3, FC3, FC1, C1, C3, AFz, Fz, F2, FC2, FCz) (fig. 3a-b). Source analysis revealed the positive cluster to be mainly driven by activity in the right postcentral sulcus, i.e., in the posterior somatosensory cortex (MNI(x,y,z) = 46 −21 42); while the later negative cluster was attributable to the posterior part of the left cingulate sulcus (MNI(x,y,z) = −15 −42 48) (fig. 3c). Finally, in line with the Self-related Facilitation account, there was also an interaction effect of HEP-FacilitationxSoundDistance on the SEP, between 161 and 206 ms after stimulation onset (cluster sum(t) = −778.31, Monte-Carlo *p* = 0.032), in a central cluster encompassing 8 channels (FC1, C1, C3, FC2, FCz, Cz, C2, C4) (fig. 3d-e). The regions most contributing to this effect were located again in the left posterior cingulate sulcus (MNI(x,y,z) = −11 −41 49), as well as in the left intraparietal sulcus (MNI(x,y,z) = −37 −44, 35) (fig. 3f). For the main effect of HEP-Facilitation, the only candidate cluster was not significant (p = 0.612).

**Figure 3.**
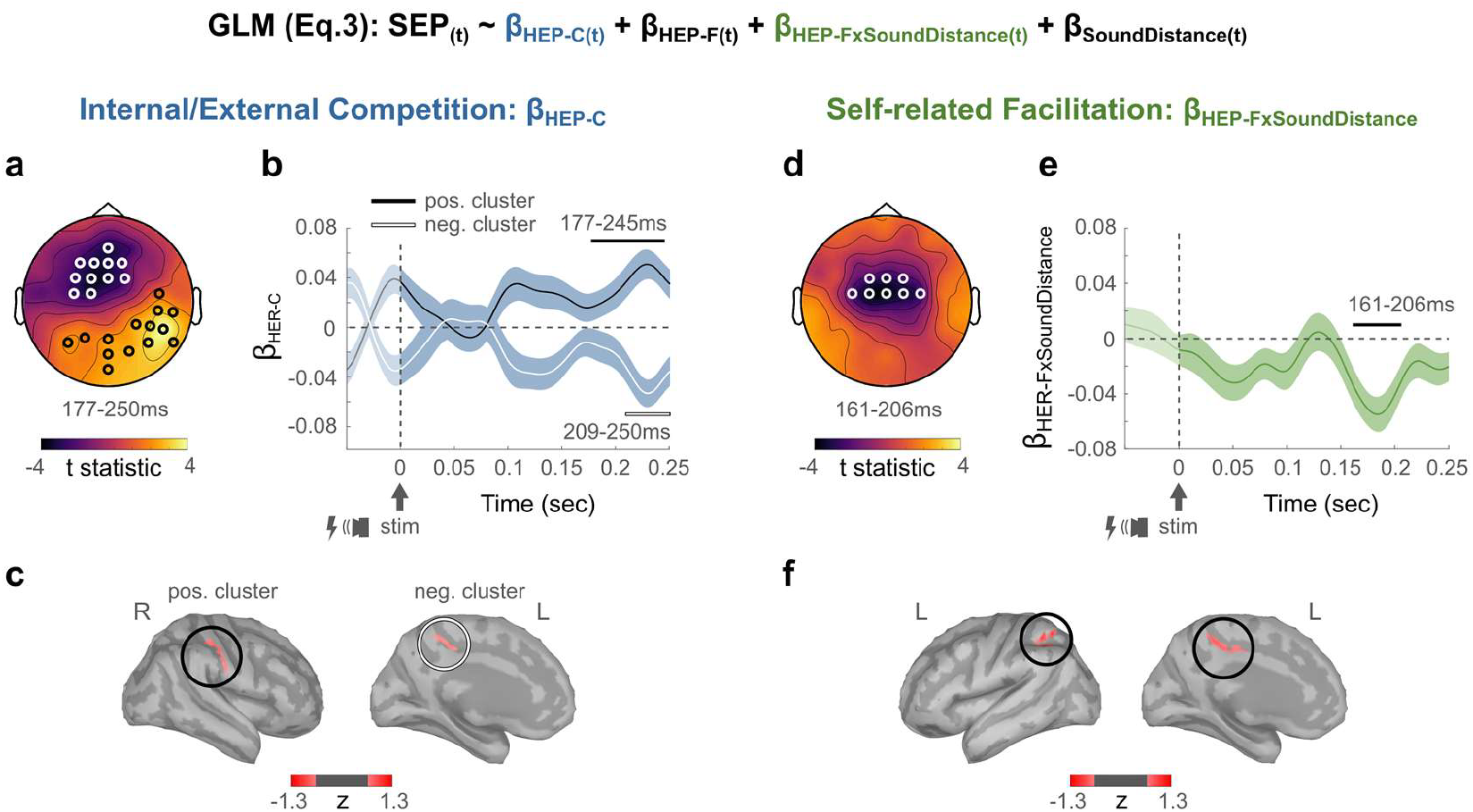
Competitive and Self-related Facilitatory effects of pre-stimulus HEPs are also reflected in audio-tactile SEPs. Pre-stimulus HEPs were averaged over the channels and timepoints showing a significant Competitive effect (HEP-Competition) and over the channels and timepoints showing a significant self-related Facilitatory effect (HEP-Facilitation) on tactile processing, and used in a GLM modelling the stimulus-evoked potentials (Eq.4). **a)** Topography of the positive and negative clusters during which there was a significant effect of HEP-Competition on the SEP, averaged over the common significant time-window (black and white bars in (b)). **b)** Timecourse of the beta estimate β_HEP-C_ averaged over the significant positive (black) and negative (white) cluster’s channels highlighted in (a). Blue areas depict standard errors. Lighter colour indicates un-tested time-window. Horizontal bars depict significant time-windows. **c)** Brain regions mostly contributing to the HEP-Competition effect on SEPs were localized in the right posterior somatosensory cortex, and in the left posterior cingulate sulcus (threshold at 50 % of max. z-value in a minimum of 20 adjacent vertices). **d)** Topography and **e)** beta estimate β_HEP-FxSoundDistance_ timecourse in the cluster during which HEP-Facilitation significantly modulated the effect of self-relevance on the SEP. **f)** Brain regions mostly contributing to the Competition effect on SEPs were localized in the left intraparietal sulcus, and in the right posterior cingulate sulcus (threshold at 60 % of max. z-value in a minimum of 20 adjacent vertices). Results for β_SoundDistance_ of the GLM (Eq.4) are depicted in supplementary fig. 2. **Abbreviations:** HEP: Heartbeat-Evoked Potential, SEP: Audio-Tactile Stimulus-Evoked Potential.

Together, these results validate and refine our previous findings. We confirmed that the observed Competition and Self-related Facilitation effects of pre-stimulus HEPs on reaction times were also reflected in the neural response to the stimulation in a corresponding pattern. Interestingly, we found that neither of the two effects affected early SEP components, and Facilitation occurred slightly earlier than Competition. Both effects modulated SEPs within the same posterior region of the cingulate sulcus, with the Facilitation effect preceding the Competition effect in this region as well. In addition, the Competition SEP effect was also driven by right somatosensory cortex, ipsilateral to the tactile stimulation, and the Facilitation SEP effect by the left intraparietal sulcus.

## Discussion

Do interoceptive signals distract us from the external world, or are they necessary components for our subjective awareness of it? By manipulating the self-relevance of a simple tactile stimulus, we designed an experiment in which the Competition and Self-related Facilitation views about interoception-exteroception interactions made distinct predictions. We found evidence in favour of both views, but linked to distinct neural substrates: HEP amplitudes before stimulation onset were negatively associated with general tactile processing, notably in somatosensory regions; but were also positively associated with self-specific processing, notably in regions of the default-mode network. Both findings were specifically related to neural processing of heartbeats rather than more general bodily states: they were time-locked to the heartbeat, and were unrelated to both the ECG, and to general arousal differences as measured by pupil diameter and heart rate. Importantly, the Competition and Self-related Facilitation effects were statistically and spatially independent from each other, suggesting coexisting, yet independent underlying mechanisms. Reinforcing this idea, only the HEP Competition effect, but not the Self-related Facilitation effect, co-varied with participants’ cardiac interoception questionnaire scores.

### 1. Competition for shared neural resources between cardiac and tactile signals in the right somatosensory cortex

Our results support the hypothesis of a competition for shared neural resources between cardiac and tactile processing, and suggest that this competition is associated with cortical processing in the somatosensory cortex: we found pre-stimulus HEPs in the somatosensory cortex to slow down reaction times and affect audio-tactile SEPs within this same region. Despite a number of methodological differences, including our use of supra-instead of near-threshold stimulation, our findings are therefore very similar to previous reports by Al and colleagues (6, 8). Our results can be interpreted within the same Internal/External Competition framework: spontaneous fluctuations in HEP amplitude reflect a trade-off in attentional resource allocation between cardiac and tactile signals. Higher HEP amplitudes would thereby indicate a state during which more resources are allocated to cardiac processing, and less resources available for the processing of incoming tactile stimuli from the external world (and vice-versa). The fact that the HEP-Competition effect positively co-varied with the propensity of participants to register cardiac sensations in everyday life reinforces the attentional competition interpretation. However, whether the Competition account can be generalised to other exteroceptive modalities beyond touch is not yet known. In fact, Park et al. (14) did not observe such competition between HEP and visual stimulus processing. Indeed, the competition we observed between tactile and cardiac processing might be entirely explained by their sharing of neural substrates within the somatosensory cortices.

Here we analysed responses to audio-tactile stimuli, but competition for neural resources should also affect unimodal tactile processing. Using tactile-only trials, we confirmed that competition was not specific to bimodal stimuli. However, the statistical effect was smaller. This is in part likely due to the much smaller number of tactile compared to audio-tactile trials in this experiment. Another possibility is that Internal/External Competition effects are generally amplified in multimodal contexts, additively affecting both simple tactile processing and subsequent multisensory integration, as recent evidence suggests that cardiac signals interact with multisensory integration involving touch (36). This remains an open empirical question.

Interestingly, we find the Competition effect within the right hemisphere, as did Al et al. (6, 8). However, in our task, this is ipsilateral to the tactile stimulation. This result might seem surprising: since tactile stimuli are predominantly processed in the contralateral hemisphere, this is where one would expect a competition for shared neural substrate. One possible explanation is that cardiac processing relies particularly on the right somatosensory cortex, consistent with proposals of a right-hemisphere specialisation of interoception (5, 37, 38). This has different consequences depending on which hand is stimulated. In our experiment, the right-hand tactile stimulus would first reach the left primary somatosensory cortex and then spread to bilateral somatosensory areas (39). Cardiac and tactile signals would therefore meet in the right hemisphere only at this later stage of tactile processing, which is when we observe interference in our data (>177 ms post-stimulation). In contrast, when the tactile stimulus is delivered to the *left* hand, both tactile and cardiac signals are processed within the right somatosensory cortex from the outset, resulting in interaction at much earlier latencies, as reported by Al and colleagues (>30 ms post-stimulation).

We did not observe any clear cardiac phase effect. Phase effects are often invoked in the context of competition between interoceptive and exteroceptive signals (7). This dissociation between HEP and phase effects suggests there might be at least two different possible competition mechanisms. Besides, phase effects are considered short-lived (one cardiac cycle by definition), and are typically interpreted with reference to predictions about the current cardiac contraction. By contrast, here, tactile processing was affected by HEPs occurring up to two seconds before stimulus onset, suggesting a mechanism operating on a much slower timescale than phase effects.

### 2. Self-related Facilitation: self-relevance as a combination of internal and external signals

In addition to Competition, however, we also found evidence for Self-related Facilitation. Indeed, spontaneous fluctuations in HEPs measured before stimulation onset predicted the ability to accurately assign self-relevance to subsequent exteroceptive stimuli: stronger HEPs predicted faster reaction times for tactile stimuli paired with a close sound, and slower reactions when paired with a far sound. This enhanced self-relevance effect was also reflected in the SEPs. Our findings thus add to the evidence according to which HEPs index self-related processing across diverse cognitive domains (for reviews, see 1, 2). Here, we provide the first direct proof that HEPs also do so during exteroception, in this instance facilitating self-relevance encoding. Because our design defined self-relevance through a contrast between peripersonal space locations of multisensory stimulation, the Self-related Facilitation account can be distinguished from a general Facilitation account, in which for instance internal and external signal processing could be jointly driven by some non-specific third factor such as arousal. One could also interpret Self-related Facilitation in attentional terms. On this view, attention could be either directed towards the self (as indexed by HEPs) and its peripersonal space, or to the rest of the external world, beyond peripersonal space. However, this type of attention would be distinct from the one involved in the Internal/External Competition effect, both in terms of scope and targets as well as neural sources and mechanisms, as detailed below.

In contrast to Competition, the Facilitation effects did not involve the somatosensory cortex, but instead higher-order integrative areas such as the medial rostral prefrontal cortex (mrPFC), the left dorsal premotor cortex and the left intraparietal sulcus. These last two areas have been linked to representations of hand-centred peri-personal space (40–42). We also found facilitating HEPs in bilateral anterior temporal regions, which are nodes of the default-mode network (43, 44), as is the mrPFC according to some authors (e.g., 45). The default-mode network is indeed involved in self-referential processing including during active tasks (e.g., 55; for a review, see 56). While self-related HEPs have previously been described in the default-mode network (9, 14, 22), they occurred at other nodes. Together, these findings suggest that HEPs might reflect a domain-general self-related signal, but that is relayed to task-specific cortical regions.

We defined self-relevance encoding as a difference between processing for stimuli occurring in near *vs* far space. While this definition cannot be completely distinguished from salience, it involves a prioritisation of stimuli with respect to body location – hence already tapping into a simple form of self. By showing that pre-stimulus HEPs facilitate this prioritisation, we further emphasize that self-relevance depends not only on characteristics of the external stimulus (in our case, sound distance), but also on the subject’s internal state – indexed here by an intrinsic neural event, the pre-stimulus HEP in premotor and default-mode network regions.

Interestingly, Self-related Facilitation was driven by slightly earlier HEP latencies and also affected earlier SEP components than Competition. These findings suggests that self-relevance encoding – and the associated facilitation by HEPs – does not correspond to slow late-stage contextualisation of sensory information. Rather, it is a relatively fast component of perceptual processing, as fast as the Competition between tactile and cardiac processing in somatosensory cortex.

### 3. The independence of Facilitation and Competition suggests multidimensionality of internal state

Crucially, Internal/External Competition and Self-related Facilitation effects were independent: they were driven by distinct neural sources, and did not correlate across participants. Of note, we did find a single region of convergence, the left posterior cingulate sulcus, where the neural response to the audio-tactile stimulus was influenced by both the Competition and the Facilitation components of the HEP. The posterior cingulate cortex is generally interpreted as a transitional zone between perception and action (48, 49), where multisensory exteroceptive (49) but also interoceptive (50) signals converge.

Our results demonstrate that HEP amplitude cannot be taken as a unitary readout of either interoception-exteroception balance or of self-related processes. Rather, the HEP needs to be understood as encoding both, and moreover in an independent manner. HEPs thus reflect at least two distinct dimensions of subjects’ internal state, each in turn differentially impacting exteroception. In other words, there is no one-to-one mapping between the neurophysiological marker HEP and a unitary concept of internal state. It follows that the specific cognitive nature of internal state cannot be inferred from scalp HEP amplitude alone.

## Conclusion

In conclusion, internal bodily signals can both distract us from external stimuli, or contribute to their perception – specifically, to the encoding of a stimulus relative to a self. We show that pre-stimulus HEPs index two distinct dimensions of a subjects’ internal state: One is linked to neural resource allocation, and produces Competition-like effects between exteroception and interoception – at least in the cardiac and tactile combination studied here. The second dimension corresponds to an integration of exteroceptive processing with the self, facilitating the discrimination between more, or less, self-relevant external events. By demonstrating the coexistence of these two dimensions, we reconcile two hitherto largely independent – and seemingly contradictory – research programmes on the relationships between interoception and exteroception. Moreover, our results highlight the multi-dimensionality of HEPs as neurophysiological markers, and thus more generally the multi-dimensionality of internal state.

## Materials and Methods

### 1. Participants

Effect size estimates for HEP effects on behaviour are not known, making formal power calculation difficult. Studies of HEP effects on perception vary in sample size between 17 (5, 14) and 40 (6, 8). We therefore aimed for a final sample size of 35, and recorded the data of 41 healthy right-handed volunteers. Four participants were excluded from data analysis due to insufficient data quality regarding EEG (less than 35 artefact-free trials per condition: 3 participants), or ECG (mean cardiac inter-beat interval under 0.6 seconds: 1 participant). Our final inferences were thus based on 37 participants (20 male, mean age ± standard deviation (SD) = 23.24 ± 2.52, range: 19-30). Experimental procedures were approved by the institutional review board of INSERM (protocol number 18-544-ter). Participants gave informed consent and received monetary compensation.

### 2. Task and Stimuli

The experimental setup and task closely followed the protocol described in Ronga et al. (31) and is illustrated in figure 1. Participants were seated in a sound-proofed room, with their head resting on a chinrest. Both arms were resting on the table in front of them, with their right hand placed close to their chest. The task was presented with Matlab 2017b (51) using the Psychtoolbox (52), running on a MacBook Air stimulation laptop.

Somatosensory stimuli were single electrical square-wave pulses of 200 *μ*s duration, applied using a constant-current stimulator (DS7A, Digitimer) to the right hand dorsum. Stimulation intensity was set to be faint, but clearly perceptible, at twice the individual detection threshold as determined by method of limits (53) (mean stimulus intensity ± SD = 3.7 ± 0.7 mA, range: 2.5 – 5.1 mA). Using such supra-threshold stimuli – instead of stimuli at threshold, as previously used in similar studies on the interactions between cardiac and tactile processing (6) – was important in order to derive meaningful reaction time measures, on which most behavioural assays of peripersonal space are based (e.g., 31, 54). Two flat 8 mm diameter surface Ag/AgCl electrodes (Neurospec AG, Stans, Switzerland) were placed either between the middle finger and index tendons, or the middle finger and ring finger tendons. After each experimental block, the electrode placement site was alternated to reduce habituation effects, and the perceptual threshold re-determined to adjust stimulation intensity.

Auditory stimuli consisted of 784 Hz pure tones of 50 ms duration, presented from one of two small FRS 8 fullrange speakers (Visaton, Haan, Germany) on the table: one speaker was placed at <10 cm from the participant’s stimulated hand, and the second was placed 130 cm away. Sounds from both speakers were physically identical: frequencies and loudness (set to ∼60 dB) were equalized using the Room EQ Wizard software and a MiniDSP EARS microphone (Audiophonics, Floirac, France). While the near sound was thus perceived as slightly louder than the far sound, we chose to use identical sound intensities at each source, to follow Ronga et al. (31)., who verified that the effect of sound distance on tactile reaction times was not affected by perceived sound loudness, but instead attributable to distance.

In each trial, either a bimodal (audio-tactile, AT) or unimodal (tactile only, T, or auditory only, A) stimulation was presented. Auditory stimuli, whether presented alone or together with a tactile stimulus, could be furthermore presented either from the near speaker (ATnear, Anear), or from the far speaker (ATfar, Afar). Subjects were instructed to respond as fast as possible to any tactile stimulus by pressing a key with their right index finger, but to ignore the sounds. Hence, A trials served as catch trials. In the bimodal condition, sounds were delivered with an (imperceptible) delay of 40 ms after tactile stimulation, following Ronga et al. (31) to account for the faster processing of auditory stimuli, and to minimise any priming of the tactile motor response by the sound.

Each trial began with a 4 to 8 seconds jittered interval, after which one of the five different stimuli (ATnear, ATfar, T, Anear, Afar) was presented. Trials ended either 0.1 seconds after response, or after a maximal response time of 1 second. During the entire task, participants were asked to fixate a dot on a screen placed behind the second speaker, and to blink only immediately following a behavioural response.

### 3. Experimental procedure

The experimental session began with the set-up of EEG and physiological recordings, as well as the electrodes for electrical stimulation. Participants then completed training for the task, consisting of an auditory perception and a task-specific training session.

The aim of the auditory training was to sufficiently familiarise participants with the near and far sounds. Participants first passively listened, and then completed at least two blocks of 20 trials each in which they had to indicate from which of the two speakers the sounds had been played. Training blocks were repeated until the error rate dropped below 20 %. At the end of the last training block, mean error rate was 4.2 ± 6.8 %.

Participants then trained for the main task. First, they were passively familiarised with each stimulus condition. They then completed a training block of 10 ATnear, 10 ATfar, 10 T, 5 Anear and 5 Afar trials, randomly intermingled. Written feedback on reaction time, missed and false alarm trials was provided at the end of the training session.

Before each experimental block of the main task, stimulation electrodes were repositioned, stimulation intensity adjusted, and the eye tracker calibrated. Each block began and ended with a 30 second baseline period, during which the participant was instructed to sit still. After the block, the screen displayed feedback about mean reaction time and hit rate. In total, participants completed five experimental blocks, each consisting of 12 randomly intermingled trials per condition, yielding a total of 60 trials per condition or 300 trials in total.

### 4. Physiological and EEG recordings

Recordings were performed in an electrically shielded room using the Biosemi ActiveTwo recording system (Biosemi, Amsterdam, The Netherlands) with a sampling rate of 1024 Hz and a DC–400 Hz bandwidth. EEG data was recorded using 64 pin-type active electrodes (Biosemi, Amsterdam, The Netherlands) with CMS and DRL placed left and right from POz respectively. Impedances were kept below 50 kΩ. The ECG was recorded using three flat-type electrodes: two under the left and right clavicles, and one used for offline re-referencing placed on the left lower abdomen. The vertical Electrooculogram (EOG) was recorded on the left eye with one electrode above the brow and one below the eye. Eye position and pupil diameter were tracked using an Eyelink 1000 system (SR Research, Canada), using a monocular recording of the right eye, with a 35 mm lens at a sampling rate of 1000 Hz. We also recorded the electrogastrogram using 8 electrodes on the abdomen, as well as respiration using a piezoelectric belt on the torso, but do not report on these measures here.

### 5. Questionnaires

Before the experimental session, participants completed online the French versions, of two online questionnaires: state anxiety from the State-Trait Anxiety Inventory (STAI) (34, French translation by 55), and the three-domain interoceptive sensations questionnaire (THISQ) (35; French translation provided by the authors), measuring interoceptive sensitivity (i.e., self-evaluated subjective interoception;, 56). We verified that all participants fell within the mean ± 3 SD of group results on all tested scales.

### 6. Preprocessing

Offline pre-processing and analysis of behavioural, physiological and neural data was done using the Fieldtrip toolbox implemented in Matlab (57) and additional custom-built Matlab code.

#### 6.1 Performance

Reaction times under 100 ms were considered as artefacts and removed from the data. For each analysed stimulus condition (T, ATnear, ATfar) separately, reaction times exceeding two SD from the participant’s mean were considered as outliers and removed from the data. Only hit trials were retained for further analysis. Reaction times were log-transformed prior to statistical modelling.

#### 6.2 ECG

We detected PQRST complexes on 1-40 Hz bandpass-filtered data of the lead II bipolar ECG derivation using a procedure based on the convolution of a template cardiac cycle with the ECG timeseries. Inter-beat intervals were calculated as the time distance between two subsequent R-peaks. Cardiac systole was defined as the interval between a cycle’s R-peak and the end of its T wave; and cardiac diastole as the interval between the end of a cycle’s T wave and the following R-peak (58). We determined end of T for each cardiac cycle using the trapezoidal area algorithm developed by Vázquez-Seisdedos et al. (59). For two subjects for which the data was too noisy, time of end of T was approximated as the maximum of the second derivative between T and T + 150 ms on their average cardiac cycle (cf. supplementary materials for more details on ECG preprocessing).

#### 6.3 EOG and Eye movements

The EOG’s bipolar derivation was computed by subtracting the electrode from under the eye from the one over the eyebrow. The eye tracker recorded saccades and pupil diameter. The pupil data was first cut into epochs of one second preceding each trial’s stimulation onset. The epochs were then automatically checked for episodes of data loss (corresponding to the camera losing track of the pupil), and artefacted epochs were excluded from further analysis.

#### 6.4 EEG

To detect bad channels, we computed the area under the curve (AUC) of each channel on the continuous, bandpass-filtered EEG data (1-45 Hz fourth order zero-phase shift forward and reverse Butterworth filter). Channels exceeding ± 3.5 SDs of the mean AUC of all channels were repaired using a weighted average of the (unfiltered) neighbouring channels. Next, the data was re-referenced to a common average of all channels. Eye movement, sharp transients, and muscle artefacts were automatically detected using FieldTrip (cf. supplementary materials). To correct potential artefacts caused by the electrical stimulation, we applied a linear interpolation on the four milliseconds following the time of stimulation. Scalp electrodes are additionally contaminated by blinks, as well as the ECG, resulting in the so-called cardiac artefact. To attenuate blink and cardiac artefacts, Independent Component Analysis was used as described in Buot et al. (32) and in the supplementary materials. Of note, this procedure has been shown to remove the influence of heart-rate and stroke volume on the heartbeat-evoked response (32). Finally, the ICA-corrected EEG data was band-pass filtered between 0.5-25 Hz using a 4th order zero-phase shift forward and reverse Butterworth filter.

### 7. Analysis

#### 7.1 Trial selection and heartbeat-evoked potential computation

Trial-by-trial HEPs were extracted as follows (illustrated in fig. 1c). For each trial, we searched for cardiac R-peaks occurring between 2000 and 600 ms prior to the stimulation onset. Only R-peaks of artefact-free cardiac cycles with a minimum inter-beat interval of 600 ms were considered. If multiple R-peaks were identified, only the R-peak closest to stimulation onset was included in the analysis. EEG data was then epoched into time windows of −30 to 600 ms time-locked to the selected R-peaks. Note that this means that HEP epochs always end before stimulation starts. HEP epochs were not baseline-corrected (as for instance in 9, 19, 21, 22), because the heartbeat is a cyclical signal – meaning that the time-window preceding an R-peak is likely to contain late responses to the previous heartbeat (1). Trials were only included in the analysis if they were free of EEG artefacts from 2000 ms preceding stimulation up to 200 ms after the response. Furthermore, only trials for which the stimulation occurred during an artefact-free cardiac cycle, and for which the participant provided a valid behavioural response, were included in the analysis. The preliminary general linear model (GLM) to compute residual reaction times was run on pooled AT and T trials, while all other analyses were run on only AT trials, which included a total of 3575 trials (mean ± SD number of trials per participant: 96.6 ± 9.2, range: 79-113; mean ± SD per condition and participant = 46.4 ± 5.1, range: 36-55).

#### 7.2 Statistical Analysis of Reaction Times and Cardiac Cycle Effects

To confirm the statistical presence of an effect of sound distance on reaction times to tactile stimuli, as well as test for an effect of cardiac phase, we applied a 2×2 repeated-measures ANOVA on log-transformed reaction times to AT trials, with two within-subject factors sound distance condition (ATnear, ATfar), and cardiac phase during stimulation onset (systole, diastole).

#### 7.3 Statistical modelling of HEPs and Behaviour

To quantify the effect of pre-stimulus HEPs on tactile processing, we used the following approach. Briefly, we first cleared each participant’s reaction times of variance attributable to cardiac parameters and order effects. Then, we modelled each participant’s trial-by-trial HEP timeseries using the residual reaction times and the sound distance condition with a general linear model (GLM). The significance of each regressor at the group level was estimated by testing the GLM’s resulting beta time series against zero across participants, using a cluster-based permutation approach. Finally, we implemented a series of control analyses. We detail each of these steps below.

##### Computing residual Reaction Times

First, we removed the effects of cardiac parameters and experimental block order on reaction times by applying a preliminary GLM (Eq.1) to the log-transformed and z-scored reaction times (RT) of pooled AT and T trials:

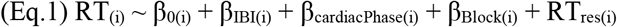

where *i* represents trial. β_0(i)_ is the intercept, β_IBI(i)_ represents the estimated contribution of the z-scored inter-beat interval of the cardiac cycle in which the trial’s stimulus occurred, β_cardiacPhase(i)_ the contribution of the phase of that cardiac cycle at stimulation onset (systole or diastole), β_Block(i)_ the contribution of the trial’s experimental block order. Finally, RT_res(i)_ are the model’s residuals, representing the part of RT_(i)_’s variance that could not be attributed to either of the three regressors. The residuals of the model were then used as the regressor RT_res_ for the remaining steps of the analysis.

For this and all subsequent GLMs, we additionally computed Variance inflation factors (VIF) between each of the regressors for each subject. All VIF values were below 1.3, indicating low multicollinearity.

##### Statistical modelling of HEPs and Reaction Times

Next, we tested for a statistical link between pre-stimulus HEPs and the speed of tactile processing on one hand, and self-relevance processing on the other hand. To this end, we modelled each participant’s trial-by-trial z-scored HEP timeseries using the following GLM (Eq.2), on AT trials:

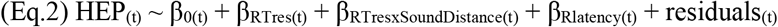

where *t* represents time. β_0(t)_ is the intercept, β_RTres(t)_ represents the estimated contribution of RT_res_, and β_RTresxSoundDistance(t)_ the contribution of the interaction between RT_res_ and the sound distance condition of the trial (near or far), i.e., the self-relevance effect. β_Rlatency(t)_ represents the contribution of the timing of the HEP’s R-peak relative to stimulation onset, and captures variance in HEPs related to heart-unspecific slow drifts in the EEG (9, 33). The resulting beta timeseries express, for each participant and each point in time and space along the HEP timeseries, the statistical link between the HEP and each regressor.

##### Testing for statistical significance at the group level

Finally, to test for significance at the group level, we compared beta timeseries against zero across participants using cluster-based permutation t-testing implemented in the Fieldtrip toolbox (60), and described in the supplementary materials. This method does not require the definition of any a priori temporal or spatial regions and intrinsically corrects for multiple comparisons in time and space. We ran 2000 cluster-based permutations with a cluster-forming threshold alpha of 0.01 and a minimum of one neighbouring electrode. To avoid contamination by any residual cardiac artefact, we only tested the time-window of 200 to 600 ms after the R-peak.

##### Control analysis: HEPs and Reaction Times within T trials

Since our main analysis was run on bimodal AT trials only, we verified that the statistical link between HEP and reaction times would also be found in unimodal tactile (T) trials, using the following GLM (Eq.3) on the z-scored HEP epochs of T trials:

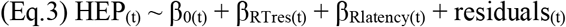

where all variables correspond to the equivalent ones in Eq.2. By experimental design, there were only half as many T trials than AT trials. We restricted this analysis to subjects for which at least 40 T trials could be included, yielding a reduced sample of 32 subjects. Furthermore, to counter this relative loss of power and since we wanted to specifically confirm the presence of the previously observed main effect of RT_res_, we also restricted the group-level cluster-based permutation to the time-points and electrodes which appeared significant in AT trials (454 to 494 ms after the R-peak, 11 electrodes: Pz, TP8, CP6, CP4, CP2, P2, P4, P6, P8, PO8 and PO4).

##### Control analysis: Statistical modelling of ECG and Reaction Times

To verify that the observed effects were truly due to a neural HEP, and not to a residual cardiac artefact, we repeated the analysis using ECG signal instead of EEG, for each of the three ECG derivations separately (32). Continuous ECG data of each derivation was segmented into epochs time-locked to R-peaks using the same procedure as for HEP computation. We then applied the GLM (Eq.2) on these ECG epochs. Cluster-based permutation testing was performed on the time-window of 200 to 600 ms after the R-peak, using a cluster-forming threshold alpha of 0.01 and no minimum of neighbouring electrodes.

##### Control analysis: Permuted heartbeats analysis

Slow waves in the EEG can confound effects attributed to pre-stimulus HEPs (1, 33). With the regressor β_Rlatency(t)_, we already separated part of the HEP’s variance due to such unspecific drifts from the variance correlating with behaviour. To further confirm that our observed effects were truly time-locked – i.e., specific – to heartbeats, we repeated the main analysis 500 times on permuted data where only the time-locking to heartbeats was broken up, and compared the resulting distribution of cluster statistics to our empirical results (14, cf. supplementary materials and for instance 19).

#### 7.4 Statistical analysis of HEPs and audio-tactile Stimulus-Evoked Potentials

In a follow-up analysis, we tested whether the effects of pre-stimulus HEPs on behaviour were also reflected in the audio-tactile stimulus-evoked potentials (SEPs). First, we computed mean HEP amplitudes over both significant clusters separately, i.e., across the timepoints and electrodes found to be significantly co-varying with either RT_res_ or RT_res_xSoundDistance, yielding two average HEP amplitude values per trial: one termed HEP-Competition and one termed HEP-Facilitation.

Then we computed the SEP for each AT trial: Continuous EEG data was segmented into epochs of −100 to 250 ms around stimulus onset, and baseline-corrected using the 100 ms before stimulus onset. For each participant, trial-by-trial z-scored SEP timeseries were then modelled at each timepoint using GLM (Eq.4):

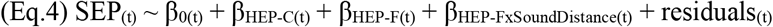

where *t* represents time, β_0(t)_ is the intercept, β_HEP-C(t)_ represents the estimated contribution of HEP-Competition, β_HEP-F(t)_ the contribution of HEP-Facilitation, and β_HEP-FxSoundDistance(t)_ the contribution of the interaction between HEP-Facilitation and sound distance on the SEP. As for the main analysis, statistical significance of each regressor at the group level was estimated using the cluster-based permutation approach, testing the full SEP time-window (0 to 250 ms after stimulus) with a first-level alpha of 0.01 and a minimum of one neighbouring electrode, using 2000 permutations.

#### 7.5 Source-level analysis

To estimate which brain regions contributed most to the observed effects, we performed source reconstruction on each of the significant regressors’ beta timeseries using the Brainstorm toolbox (61) with a default anatomy (ICBM152) and no noise modelling. Prior to averaging, individual source maps were z-scored using an automatically detected “baseline” period in which beta amplitude and variance was the smallest across subjects. This step was necessary because cardiac-related data lack a proper baseline. Details of the source reconstruction procedure are described in the supplementary materials. The z-scored source maps were then averaged across subjects and respective timepoints, separately for each significant cluster, revealing the cortical structures most contributing to each effect.

#### 7.6 Correlations with the Competition and Facilitation effects

Finally, we tested whether Competition and Facilitation effects were correlated with each other within subjects, and further explored whether they were modulated by either subjects’ trait anxiety or by their cardiac interoceptive sensibility. Individual Competition and Facilitation effects were quantified by averaging each participants’ beta values across group-level significant time windows and electrodes. We then computed robust correlations between effects, and between effects and questionnaire scores. False-Discovery Rate (FDR) correction for multiple comparisons was applied for each questionnaire separately (i.e., for THISQ across the two subscales of interest and the two beta values, for STAI across the score and the two beta values). Evidence in favour of correlations was quantified by computing Bayes Factors BF10, using the Matlab toolbox bayesFactor (62). Values between 0.1 and 0.3 indicate moderate evidence in favour of the null hypothesis H0, while values between 3 and 10 indicate moderate evidence in favour of H1.

## Acknowledgments

We thank Véronique Marchand-Pauvert for generously lending us the electrical stimulation device, and Francesca Garbarini for her helpful advice regarding the paradigm. This research was supported funded by Agence Nationale pour la Recherche (ANR-17-EURE-0017 and ANR-10-IDEX-0001-02), as well as a senior fellowship of the Canadian Institute For Advance Research (CIFAR) program in Brain, Mind and Consciousness to C.T.-B. P.H. was supported by a Reimar-Lüst Preis of the Humboldt Foundation. A CC-BY-NC public copyright license has been applied by the authors to the present document and will be applied to all subsequent versions up to the Author Accepted Manuscript arising from this submission, in accordance with the grant’s open access conditions.

## Data & Code availability

The custom code is available online (https://osf.io/yzmx4/?view_only=802fd7c48dd74d5fa1d6957d105f4294). Data supporting main results and figures are shared in the same repository. To comply with EU laws, individual data can only be shared once an institutional data sharing agreement with Inserm has been signed.

## Author Contributions

M.L.: conceptualisation, methodology, formal analysis, writing original draft & editing. P.H.: conceptualisation, methodology, writing – review & editing, supervision. C.T.-B.: conceptualisation, methodology, formal analysis, data curation, writing original draft & editing, funding acquisition, supervision, project administration.

