## Supplementary Materials for "Interoception vs. Exteroception: Cardiac interoception competes with tactile perception, yet also facilitates self-relevance encoding"

**This PDF file includes:**

Supporting text
Figures S1 to S2
SI References

**Supporting Information Text**

**Supplementary Details on Methods**

**1. ECG preprocessing: Detection of PQRST complexes and end of the T wave**

First, we computed the lead II bipolar derivation, between the right clavicle ECG electrode and the reference electrode of the abdomen. We then detected R peaks: After band-pass filtering between 1 and 40 Hz (windowed-sinc FIR filter), a template cardiac cycle was computed and convolved with the ECG time series. R-peaks were detected on the normalized (between 0 and 1) resulting convolution, as a correlation exceeding a threshold of 0.6. The other components of each cycle's PQRS complex were detected as local minima or maxima in fixed time-windows relative to the R peak latency. For each subject, we visually inspected the margins of the distributions of R-to-R, R-to-T and P-to-R time intervals, and manually corrected the automatically detected peaks when necessary and possible. Remaining outlier cycles in R-to-T and P-to-R intervals, defined as an interval exceeding 3 SD from the mean or one interquartile range, respectively, were automatically removed from further analysis.

The ends of each cardiac cycle's T wave were detected using the trapezoidal area algorithm developed by Vázquez-Seisdedos et al. (1). As this method operates on each cardiac cycle individually, it accounts for between- and within-subject variability in the length of cardiac phases; but is also more sensitive to noise. For cardiac cycles in which the data was too noisy to apply the trapezoid method, we estimated the time of end of T based on the mean of that block's average end of T time (on average for 0.09 ± 0.45 % of cycles). Subjects for which the trapezoid method yielded estimations with a variance exceeding 2 SD from the group's mean were deemed too noisy for this method (2 subjects). For them, time of end of T was approximated as the maximum of the second derivative between T and T+150ms on their average cardiac cycle.

**2. EEG preprocessing: Detection and attenuation of artefacts**

**2.1 Automatic artefact detection**

Eye movement artefacts were detected as saccades exceeding 3 degrees of amplitude. To detect jump artefacts, EEG data was median filtered (24th order) and z-scored; sharp transient artefacts were defined as segments during which a channel's z-score exceeded a cut-off value of 20, with 100 ms padding. Channels containing more than 10 such shifts in one block were repaired using neighbouring channels as described above. To detect muscle artefacts, EEG data was band-pass filtered (110-140 Hz, 8th order zero-phase forward and reverse Butterworth filter), Hilbert transformed, and z-scored; muscle artefacts were defined as segments during which the signal across channels exceeded a cumulated z-score of 10, with 100 ms padding.

**2.2 Attenuation of the blink and cardiac artefacts with ICA**

Scalp electrodes detect not only neural activity but are also contaminated by blinks, as well as by the electrocardiogram, resulting in the so-called cardiac artefact in the EEG signal. To attenuate blink and cardiac artefacts, Independent Component Analysis (ICA) was used as described in Buot et al. (2). For blink correction, the re-referenced and stimulus-artefact-free EEG and EOG data were high-pass filtered at 0.5 Hz using a fourth order zero-phase shift forward and reverse Butterworth filter, and epoched into segments free of artefacts but containing at least one blink event). Data was then decomposed into independent components using the ft_componentanalysis fieldtrip function. Next, pairwise phase consistency (PPC) was computed between the frequency decompositions of the ICA components and the EOG data in the 0-15 Hz range. Up to three components were identified as blink-related if they exceeded more than three standard deviations from the mean PPC. Identified components were removed from the continuous re-referenced and stimulus-artefact-free EEG data using the ft_rejectcomponent function.

For the attenuation of cardiac artefacts, blink-corrected EEG and re-referenced ECG data were high-pass filtered at 0.5 Hz using a 4th order zero-phase shift forward and reverse Butterworth filter, and epoched into segments of 400 ms centred on the R-peaks. As for the blink ICA, artefact-free EEG data segments were then decomposed into independent components, and PPC computed between each independent component and the ECG signal in the 0-25 Hz range. Up to three components exceeding more than three standard deviations from the mean PPC were identified as related to cardiac events, and removed from the blink-corrected EEG data.

### 3. Analysis

#### 3.1 Testing for statistical significance at the group level: Cluster-based permutation testing

To test for significance at the group level after fitting GLMs to each participants' EEG data, we compared the resulting beta timeseries against zero across participants using cluster-based permutation t-testing implemented in the Fieldtrip toolbox (3). For each beta timeseries, two-tailed paired *t*-tests are used to compare each sample at each electrode to zero. Individual samples with a *t*-value exceeding the cluster-forming statistical threshold are clustered together based on temporal and spatial adjacency, and considered as candidates given a set minimum number of neighbouring electrodes. Candidate clusters are characterized by the sum of the *t* values of their individual samples sum(*t*). The statistical significance of candidate clusters is then determined using the Monte Carlo method: Condition labels (beta vs. null) are randomized across participants 2000 times. For each randomization, the *t*-testing is repeated and the largest (resp. smallest) sum(*t*) of the observed clusters is selected to generate the distribution of largest (resp. smallest) sum(*t*) under the null hypothesis. The Monte Carlo *p* value, expressing the likelihood of observing the original cluster-level test statistic under the null hypothesis, then corresponds to the proportion of sum(*t*) that exceed (resp. is inferior to) the original sum(*t*). We ran cluster-based permutations between each regressor's beta timeseries and zero using the clustering procedure with a threshold alpha of 0.01 and a minimum of one neighbouring electrode. To avoid contamination by any residual cardiac artefact, we only tested the time-window of 200 to 600 ms after the R-peak.

#### 3.2 Control: Checking heartbeat-specificity using Permuted heartbeats analysis

The permuted heartbeats analysis, as previously implemented for instance by Park et al. (4) and Engelen et al. (5), is used to verify that observed effects are truly timelocked – i.e., specific to – hearbeats, instead of being attributable to unspecific slow drifts in the data affecting both HEPs and any other pre-stimulus neural measure. For each participant, we permuted heartbeat timings with respect to stimulus onsets: the original timings of the R-peaks used in the HEP analysis were reassigned randomly, such that the R-peak timing of trial *i* was reassigned to trial *j*. Continuous EEG data was epoched into permuted "HEPs" around the new permuted "R peaks". The GLM (Eq.2) was then re-run on these new epochs, the same group-level cluster-based permutation procedure was applied, and the largest (resp. smallest) sum(*t*) value selected. If no candidate cluster was identified in the permuted data, a sum(*t*) value of 0 was assigned to the permutation round. The procedure was repeated for a total of 500 permutations. Finally, the cluster statistic of each of the observed clusters in the original HEP analysis was compared to the respective distribution of permuted cluster statistics to derive the Monte-Carlo *p*, reflecting the probability of the result being not truly locked to heartbeats.

#### 3.3 Estimating neural sources of observed effects on HEP data

To estimate which brain regions contributed most to the observed effects, we performed source reconstruction on each of the significant regressors' beta timeseries using the Brainstorm toolbox (6) with a default anatomy (ICBM152) and the Biosemi 64 10-10 layout. A 3-shell sphere was created for each participant. No noise modelling was performed. Sources of each participant's beta timeseries were computed using default parameters (minimum norm imaging, current density map measure, constrained normal to cortex source model, depth weighting order 0.5 with maximal amount 10, regularization parameter signal-to-noise ratio of 3 and output mode inverse kernel only); noise covariance was set to diagonal.

Baseline normalization of source maps prior to group-level averaging is recommended by Brainstorm to avoid a bias of estimations toward superficial sources when using minimum norm imaging. However, HEPs being cyclical, the beta timeseries of the GLM (Eq.2) on HEPs lack a pre-event baseline period. Instead, we determined a relatively silent period of 100 ms for each of the two beta timeseries separately, which we then used to z-score source maps. For each possible 100 ms time window between -30 and 600 ms after the R peak, we computed the standard deviation and the average absolute amplitude across electrodes. We then selected the time window for which signal amplitude and variance was smallest. The so selected “baseline” windows were 42 to 142 ms after the R peak for $\beta_{RTres}$, and 136 to 236 ms for $\beta_{RTresxSoundDistance}$. The windows did not include any significant timepoint. Each participant’s estimated source map was z-scored using the mean and standard deviation of this window. For the beta timeseries of GLM (Eq.3) on SEPs, we z-scored using the 100 ms baseline period preceding stimulation.

The z-scored source maps were then averaged across subjects and respective timepoints, separately for each significant cluster, revealing the cortical structures most contributing to each effect.

### Supplementary Results

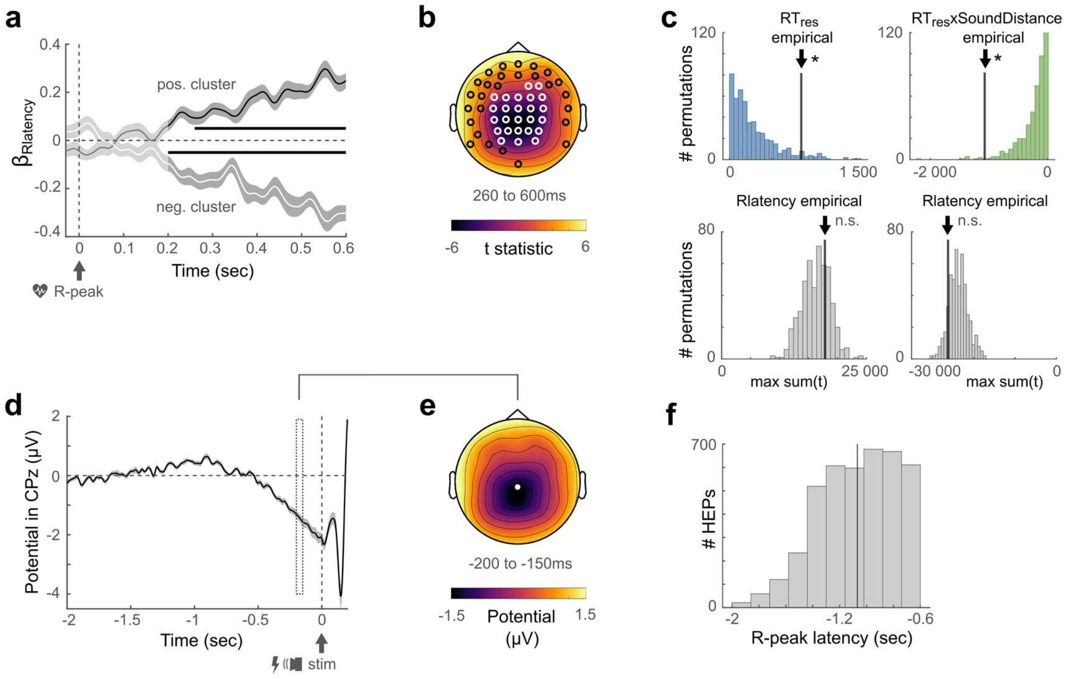


**Fig. S1. Timing of R-peak (Rlatency) relative to stimulus onset captures variance related to a non-specific slow drift. a)** Timecourse of the positive (black) and negative (white) cluster's estimate βRlatency of GLM (Eq.2), averaged across respective significant channels highlighted in (b). Shaded areas depict standard errors. Lighter colour indicates un-tested time-window. Significant time-windows are denoted by black horizontal bars. **b)** Topography across the common significant time-window. **c)** Results of the permutation control analysis, indicating that the effects of pre-stimulus HEPs on $RT_{res}$ (Competition) and RTresxSoundDistance (Facilitation) were truly time-locked and hence specific to the cardiac R-peak as they do not lie within 95% of the distribution of maximal sum(t) statistics obtained by permuting R-peak timings. As expected, the positive and negative effects of the regressor Rlatency, meant to capture variance in HEPs related to slow drifts not locked to heartbeats, do not survive this permutation control analysis. **d)** Group-averaged timecourse at channel CPz (highlighted in white in (e)) before stimulation onset, showing a slow negative drift up to stimulation. Shaded area depicts standard error. **e)** Topography of negative pre-stimulus potential; note the similarity to (b). **f)** Distribution of R-peak timings relative to stimulus onset of HEPs included in the analysis. Vertical bar marks the mean. Most HEPs fall within the time-window of the negative drift, explaining the Rlatency effect. **Abbreviations:** HEP: Heartbeat-Evoked Potential, Rlatency: Timing of cardiac R-peak relative to stimulus onset.

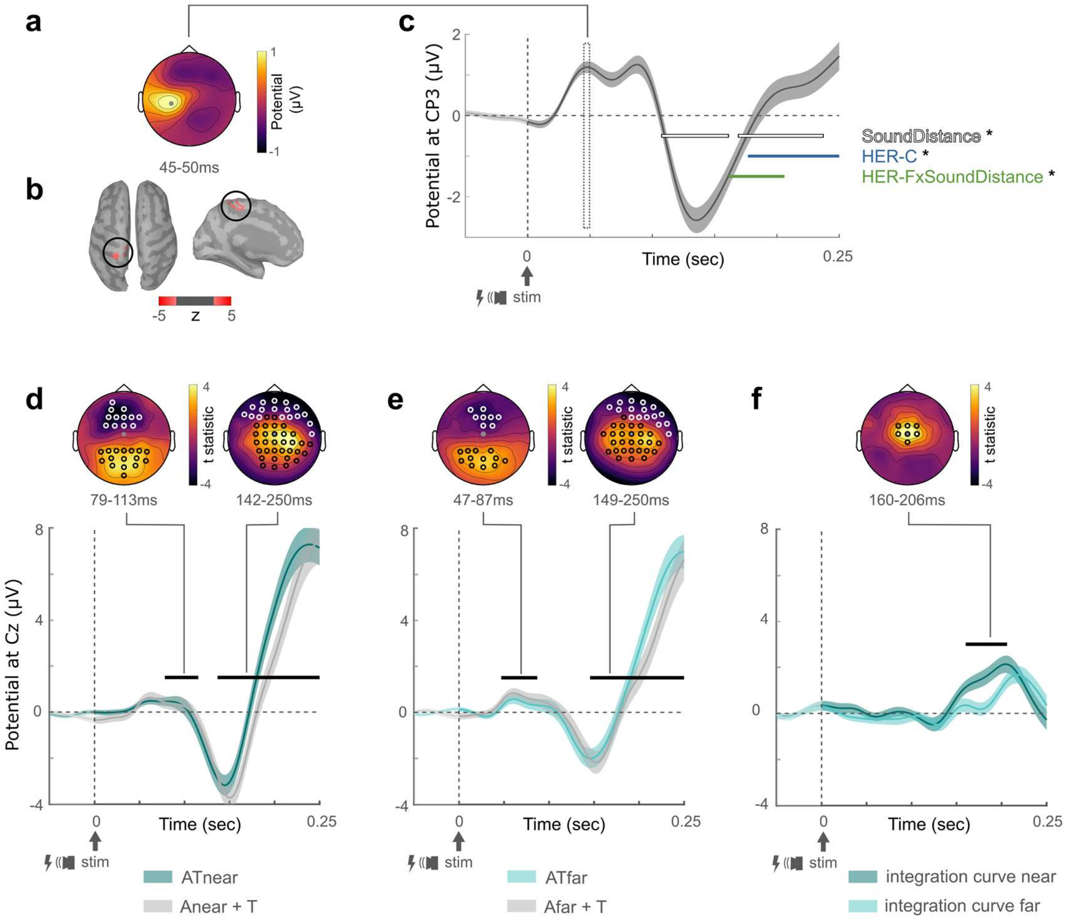


**Fig. S2. Effects of sound distance on the audio-tactile Stimulus-Evoked Potential.** Upper panels: Group-averaged SEP of AT trials. **a)** Topography of the P50 component, depicting the typical positive potential over the somatosensory cortex contralateral to the stimulated hand. **b)** Estimated sources of the P50 (threshold at 66 % of max. z-value in a minimum of 20 adjacent vertices). **c)** Timecourse of the SEP at electrode CP3 (highlighted in grey in (a)). Shaded area depicts standard error. Lighter colour indicates un-tested time-window. Time-windows during which sound distance and pre-stimulus HEPs significantly modulated the SEP are marked by horizontal bars. **d)** Results of the permutation-based cluster analysis contrasting the SEP of ATnear trials (turquoise) to the sum of SEPs of Anear and T trials (grey), at the channel Cz (grey filled dot in the topographies). Time-windows during which the two curves significantly differed from each other are marked by the black bars. Channels at which the curves significantly differed from each other are highlighted in white on the respective topographies. Significant differences between the curves indicate a supra- or sub-additive response to bimodal trials compared to summed unimodal trials, i.e., the signature of true multisensory integration (Ronga et al., 2021). **e)** Results of the contrast between SEP of ATfar trials (turquoise) and the summed SEPs of Afar and T trials (grey). **f)** To verify that multisensory integration was stronger within peripersonal space, we computed integration curves for ATnear and for ATfar conditions separately by subtracting the summed unimodal responses (Anear+T, Afar+T) from the respective bimodal responses (ATnear, ATfar). The near integration curve (dark turquoise) was significantly higher than the far integration curve (light turquoise) over the time-window depicted by the black bar and channels highlighted in the topography above, indicating stronger multisensory integration in near space. **Abbreviations:** SEP: Audio-Tactile Stimulus-Evoked Potential, HEP: Heartbeat-Evoked Potential, HEP-C: HEP-Competition, HEP-F: HEP-Facilitation, AT: Audio-Tactile, A: Audio, T: Tactile.
